# A repurposed, blood gene signature is associated with poor outcomes in SARS-CoV-2

**DOI:** 10.1101/2020.11.21.392670

**Authors:** Brenda M. Juan-Guardela, Jiehuan Sun, Tong Zhang, Bing Xu, Gaetane Michaud, Jose D. Herazo-Maya

## Abstract

Poor outcomes after SARS-CoV-2 infection are difficult to predict. Survivors may develop pulmonary fibrosis. We previously identified a 52-gene signature in peripheral blood, predictive of mortality in Idiopathic Pulmonary Fibrosis. In this study, we analyzed this signature in SARS-CoV-2 infected individuals and identified genomic risk profiles with significant differences in outcomes. Analysis of single cell expression data shows that monocytes, red blood cells, neutrophils and dendritic cells are the cellular source of the high risk gene signature.

## Main

SARS-CoV-2 infection is associated with increased mortality and poor outcome in individuals at risk. Unfortunately, it is difficult to predict which patient will deteriorate, require ICU admission and mechanical ventilation. While patients who receive invasive mechanical ventilation are more likely to be males, obese, and to have elevated values of liver-function tests and inflammatory markers (ferritin, d-dimer, C-reactive protein, and procalcitonin)^1^, reliable, peripheral blood biomarkers of disease severity and poor outcomes are still necessary to better triage patients and improve utilization of resources. Given the fact that pulmonary fibrosis can be seen in certain SARS-CoV-2 survivors^2^, we hypothesized that a 52-gene signature, previously shown to predict mortality in Idiopathic Pulmonary Fibrosis (IPF)^3,4^ could be repurposed to predict poor outcomes in SARS-CoV-2 infection.

To test our hypothesis, we analyzed peripheral blood gene expression levels of 50 genes of the 52-gene signature from SARS-CoV-2 infected subjects from a discovery (11 subjects)^5^ and validation (100 subjects)^6^ cohorts. A cellular-source cohort (7 subjects)^7^ was used to identify the cellular source of the 50-gene signature. Two non-coding RNAs were not found in some of the datasets and excluded from the original signature for consistency. This signature consisted of seven genes with increased expression and 43 genes with decreased expression. The scoring algorithm of molecular phenotypes (SAMS)^4^ (see methods section) was used to identify high and low genomic risk profiles based on the 50-gene signature in each cohort (Figure 1A and 1B). Subjects with a high risk genomic profile were more likely to have severe disease (P=0.045) in the discovery cohort. In the validation cohort, subjects with a high risk genomic profile were significantly older with higher Apache II and Charlson Disease Severity Index scores. They were more likely to be admitted to ICU, to be on mechanical ventilation, to have shorter ventilator-free days, hospital-free days, higher lactate, CRP and d-dimer levels (see Table 1).

**Table 1.**
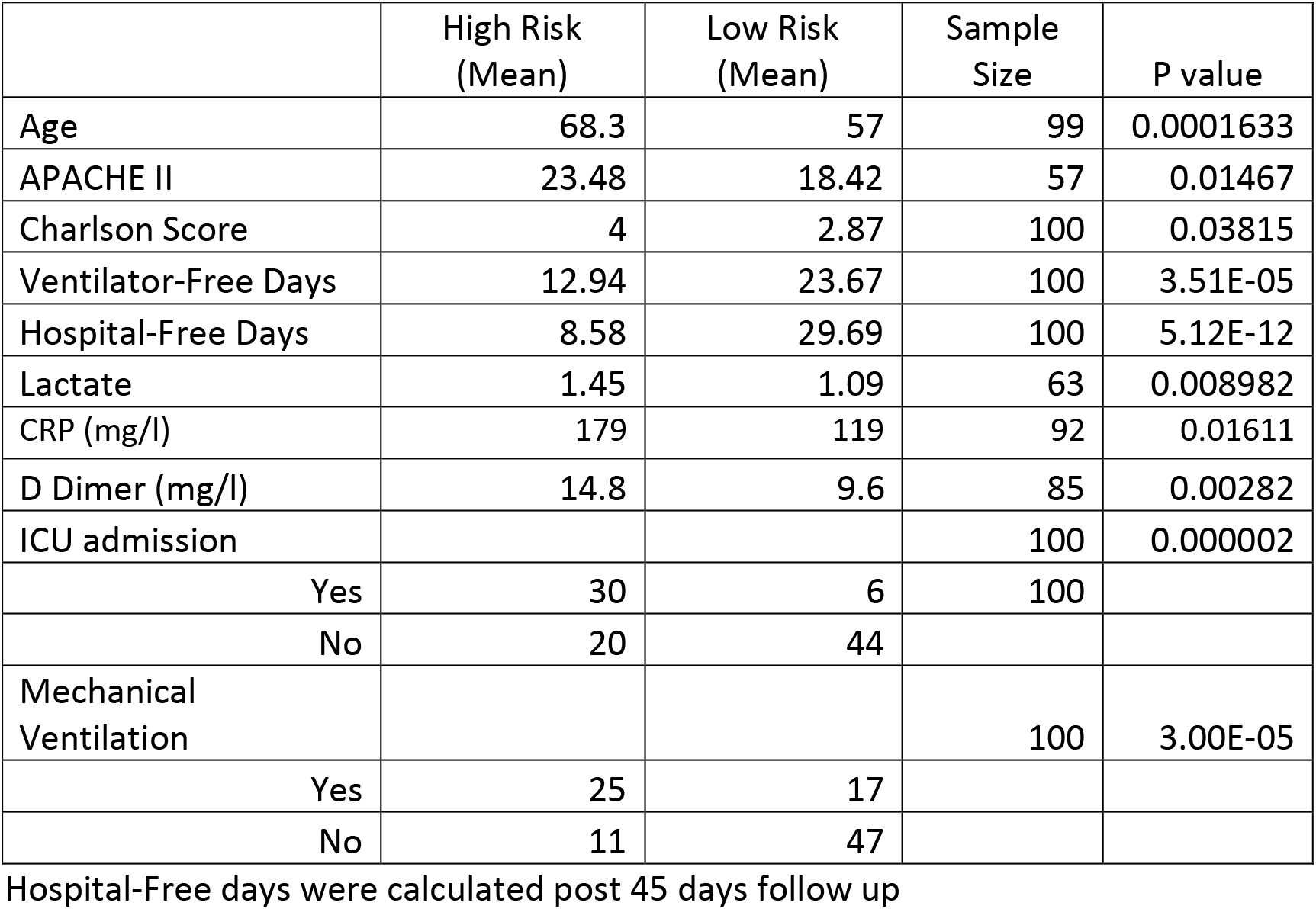
Validation cohort.

|  | High Risk<br>(Mean) | Low Risk<br>(Mean) | Sample<br>Size | P value |
| --- | --- | --- | --- | --- |
| Age | 68.3 | 57 | 99 | 0.0001633 |
| APACHE II | 23.48 | 18.42 | 57 | 0.01467 |
| Charlson Score | 4 | 2.87 | 100 | 0.03815 |
| Ventilator-Free Days | 12.94 | 23.67 | 100 | 3.51E-05 |
| Hospital-Free Days | 8.58 | 29.69 | 100 | 5.12E-12 |
| Lactate | 1.45 | 1.09 | 63 | 0.008982 |
| CRP (mg/l) | 179 | 119 | 92 | 0.01611 |
| D Dimer (mg/l) | 14.8 | 9.6 | 85 | 0.00282 |
| ICU admission |  |  | 100 | 0.000002 |
| Yes | 30 | 6 | 100 |  |
| No | 20 | 44 |  |  |
| Mechanical<br>Ventilation |  |  | 100 | 3.00E-05 |
| Yes | 25 | 17 |  |  |
| No | 11 | 47 |  |  |

Hospital-Free days were calculated post 45 days follow up

**Figure 1.**
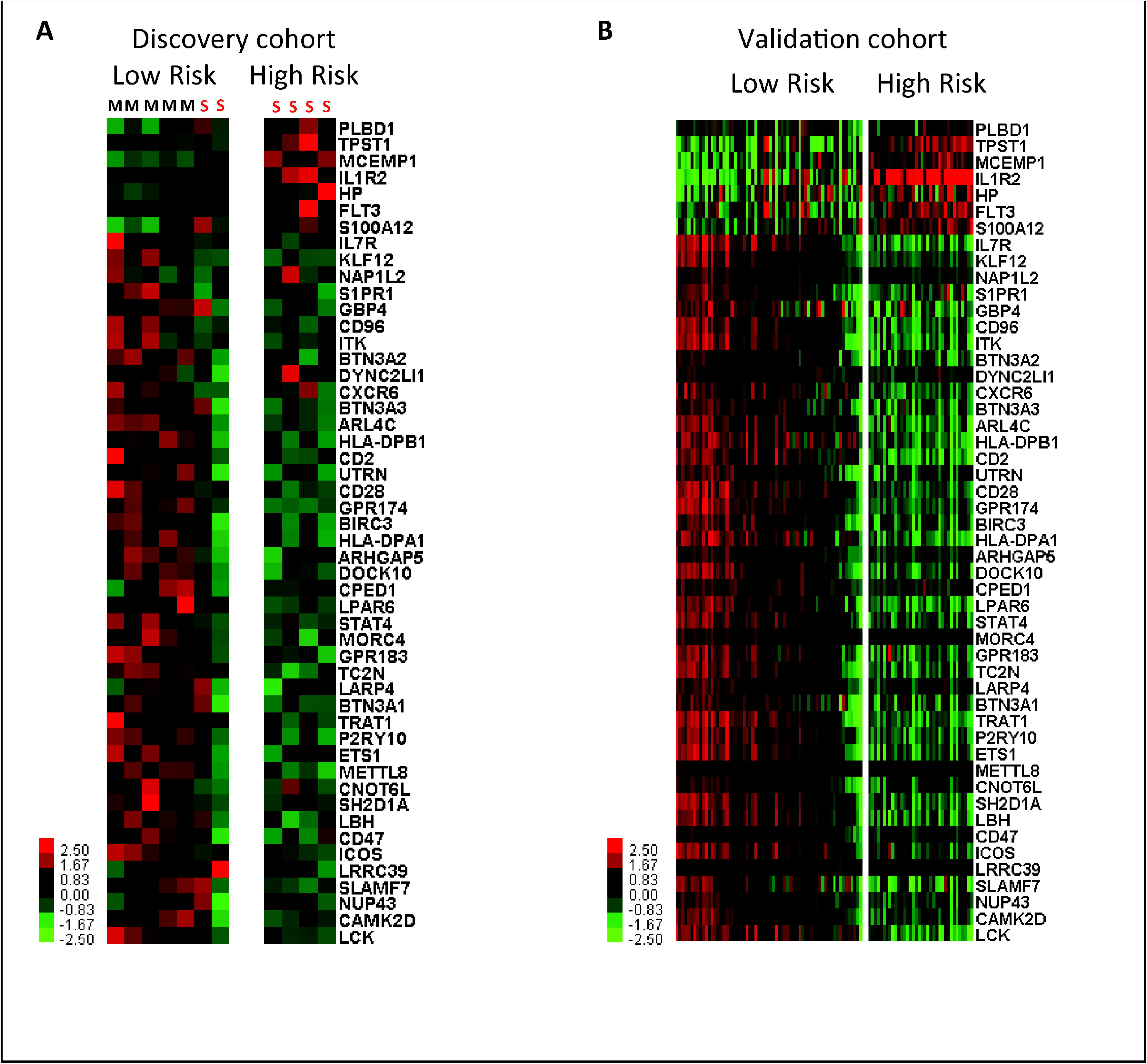
Genomic risk profiles based on the 50-gene signature are predictive of poor outcomes in SARS-CoV-2. Clustering of SARS-CoV-2 infected individuals based on genomic risk profiles (high vs low) derived from the 50-gene signature using SAMS in discovery (A) and validation cohorts (B). Every column represents a subject and every row represents a gene. Log-based two color scale is shown next to heatmaps; red denotes increase expression over the geometric mean of samples and green, decrease. S=SARS-CoV-2 severe disease. M=SARS-CoV-2 mild disease

To identify the cellular source of the 50-gene signature, we conducted both cell-type and subject-level analysis using eight, single-cell data measurements from seven SARS-CoV-2 infected subjects. For the subject-level analysis, SAMS classified five low risk and three high risk profiles using the average expression levels of each gene across all cell types. We then compared the proportions of different cell types between high and low risk profiles and identified that subjects with high risk profiles had higher proportion of CD14^+^ monocytes, red blood cells, CD16^+^ monocytes, neutrophils and dendritic cells. Subjects with low risk profiles had higher proportions of CD4 and CD8 T cells, natural killer, B cells and immunoglobulin-producing plasmablasts (Figure 2A). To validate these findings, we conducted cell-type-level analysis. Specifically, we estimated the average expression levels of each gene, for each cell type producing 155 cell-type-specific expression profiles, among which SAMS identified 46 high risk profiles and 109 low risk profiles (Figure 2B). Cell types with high risk profiles included dendritic cells, CD16^+^ monocytes, neutrophils, eosinophils, CD14^+^ monocytes, red blood cells and plasmacytoid dendritic cells. Cell types with low risk profiles were mostly CD4 and CD8 T cells, natural killer, B cells and immunoglobulin producing plasmablasts (Figure 2C) further confirming that these cells are likely the cellular source of the risk profiles, based on the 50-gene signature.

**Figure 2:**
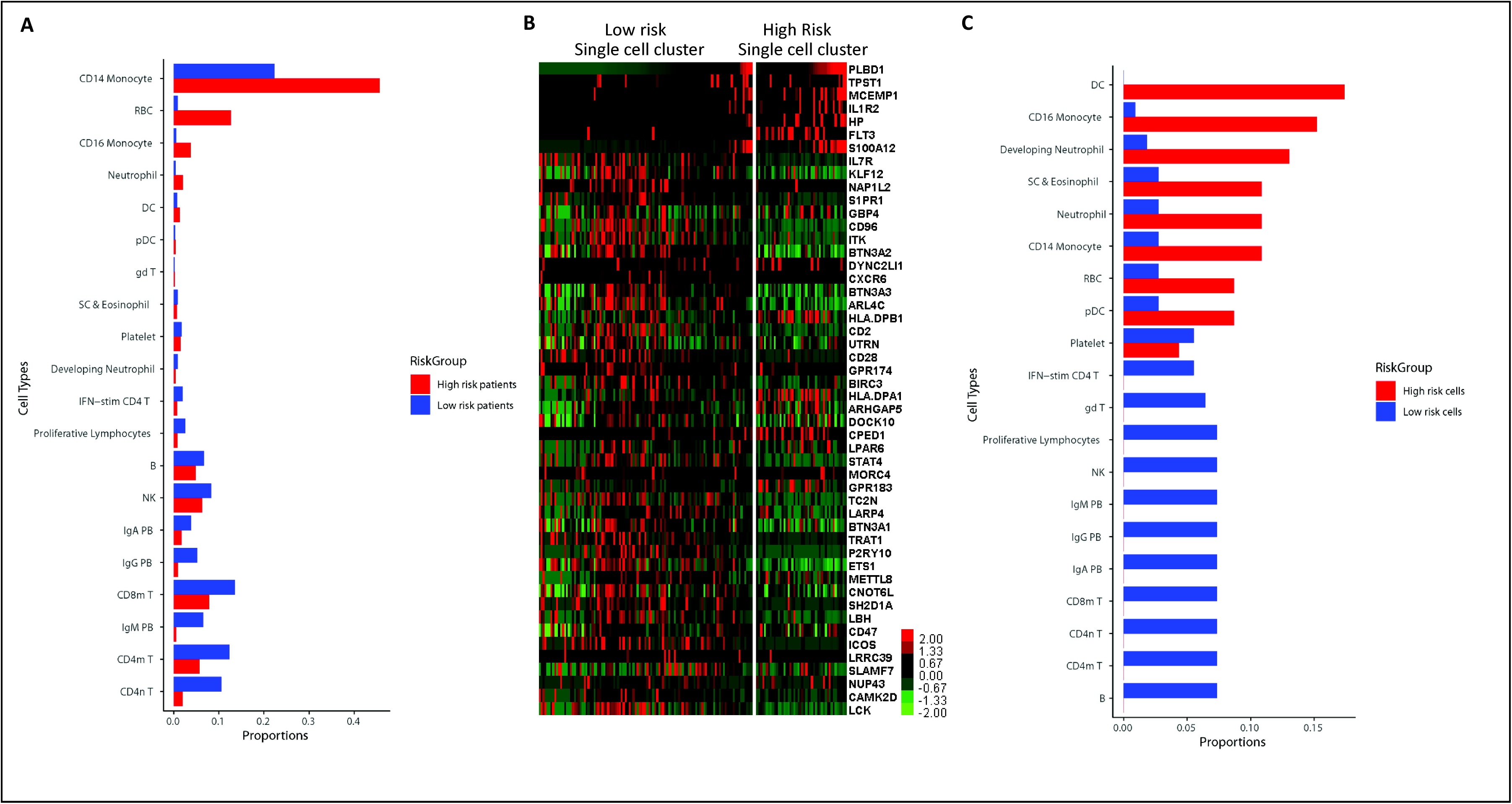
(A) Analysis of subject-level, single cell expression of the 50-gene signature from the cellular cohort demonstrates differences in cell proportions in high versus low risk individuals. Y axis represents cell types and X axis represents cell proportions. B: B Cell, CD4m T: Memory CD4 T Cell, CD4n T: Naive CD4 T Cell, CD8m T: Memory CD8 T Cell, DC: Conventional Dendritic Cell, gd T: Gamma Delta T cells, IFN-stim CD4 T: Interferon-stimulated CD4 T cell, IgA PB: IgA (Immunoglobulin-A) Plasmablast, IgG PB: IgG (Immunoglobulin-G) Plasmablast, IgM PB: IgM (Immunoglobulin-M) Plasmablast, NK: Natural Killer Cell, pDC: Plasmacytoid Dendritic Cell, RBC: Red Blood Cell, SC & Eosinophil: Stem Cells and Eosinophil. (B) Cell-type-level analysis. Heatmap shows cell types with low versus high risk expression profiles based on the 50-gene signature. Every column represents a single cell and every row represents a gene. Log-based two color scale is shown next to heatmaps; red denotes increase expression over the geometric mean of samples and green, decrease. (C) Cell-type-level expression analysis of the 50-gene signature showed that the high-risk cell cluster mainly consisted of Dendritic cells, CD16+ monocytes, Neutrophils, Eosinophils, CD14+ monocytes, Red Blood Cells and pDC. Low risk cell clusters mainly consisted of CD4 and CD8 T cells, Natural Killer cells and Immunoglobulin producing plasmablasts.

In summary, we repurposed a 50-gene signature previously shown to predict mortality in IPF and demonstrated that it is predictive of poor outcomes in SARS-CoV-2 infected subjects from two independent cohorts. The biomarker implications of this discovery are significant since the identification of SARS-Cov-2 genomic risk profiles based on the 50-gene signature, in addition to other biomarker and clinical variables, can aid with healthcare utilization such as triage of patients to most appropriate location (home, ward, ICU), reduce cost of inappropriate hospitalization, decrease personal protective equipment use and allow for proper allocation of limited resources. It may also serve for clinical trial purposes since high risk subjects may respond differently to prolonged courses of Remdesivir, higher doses of dexamethasone or new therapies.

The analysis of single cell data in SARS-CoV-2 infected subjects uncovered the cellular source of the 50-gene signature and pointed at CD14^+^ monocytes and neutrophils as critical cell types differentiating high versus low risk individuals (Figure 2A). Previous reports have shown that severe COVID-19 is marked by the occurrence of neutrophil precursors and increased circulating levels of CD14^+^ monocytes with high expression of alarmins *S100A8/9/12 and* low expression of *HLA-DR*^*8*^. Our single cell analysis also confirmed the presence of increased number of neutrophil precursors, mature neutrophils and CD14^+^ monocytes with high expression of S100A12 and low expression of HLA-DPB1 and HLA-DBP1 as part of the high risk genomic profile (Figure 2A, 2B and 2C). We have previously shown increased circulating levels of CD14^+^ monocytes in IPF patients with a high risk genomic profile based on the 52-gene signature^9^. We also showed that increase circulating levels of CD14^+^ monocytes were predictive of transplant-free survival and mortality in IPF and other fibrotic diseases such as, scleroderma-associated interstitial lung disease (SSc-ILD), hypertrophic cardiomyopathy and myelofibrosis^9^. Our single cell analysis also showed increased proportion of CD4 and CD8 T cells and Immunoglobulin producing plasmablast in individuals with a low risk profile, suggesting that strong T cell^10^ and distinct antibody responses^11^ may be responsible for milder COVID-19 infection seen in individuals with a low risk genomic profile. Future studies correlating genomic risk profiles with immunophenotyping will be required to better understand the gene expression changes seen in subjects with a low risk genomic profile.

Despite the reproducibility of our biomarker results we need to acknowledge some limitations of our study. First, given the experimental differences in discovery and validation cohorts, we need to rely on gene expression normalization within cohorts to identify genomic risk profiles based on the 50-gene signature. Second, this study was retrospective and our results will need further validation using the same gene expression platform, multiple and larger patient cohorts with samples collected prospectively. Lastly, future studies will be needed to address whether the 50-gene signature can predict mortality in SARS-CoV-2 infected individuals.

In conclusion, our study shows that a 50-gene signature in peripheral blood, previously shown to predict mortality in IPF, is able to distinguish two genomic risk profiles with significant differences in outcomes in patients infected with SARS-CoV-2 virus. It also provided evidence regarding the cellular source of the signature. A blood test based on the 50-gene signature should be further developed and validated to be used in clinical practice for risk stratification in COVID-19 infection.

## Methods

Gene expression data of SARS-CoV-2 infected individuals were analyzed using discovery (11 subjects with mild or severe SARS-CoV-2, single cell gene expression, GEO: GSE149689), validation (100 subjects, bulk cell RNA-seq gene expression, GEO Accession: GSE157103^6^) and cellular source (7 subjects, single cell gene expression, GEO accession: GSE150728^7^) cohorts. All analyses were performed in R software (version 4.0.2)^12^. For the single cell dataset in the discovery cohort, we used R package “Seurat” to preprocess the feature-barcode matrices obtained from the GEO database. Specifically, we discarded cells that expressed less than 200 genes or expressed mitochondrial genes in >15% of their total gene expression, and discarded genes that expressed in less than 10 cells. NormalizeData function was used to obtain normalized gene expression levels. Subject-level expression profile was estimated using the average expression level across all cells. For the RNA-seq data in the validation cohort, we obtained the Transcripts Per Million (TPM) matrix from the GEO database and used log(1+TPM) to normalize the gene expression levels. For the single cell data in the cellular source cohort, we used the preprocessed and normalized data provided in the published paper^8^.

To identify high and low genomic risk profiles for subjects and cells in each one of these cohorts, we used the Scoring Algorithm for Molecular Subphenotypes (SAMS). SAMS is a classification algorithm developed to identify molecular subphenotypes based on the expression of a predefined set of increased and decreased genes in a given patient. The up and down score of SAMS are calculated using the product of two variables: the proportion of genes expected to be increased or decreased per subject and their normalized expression levels. The calulation steps of SAMS have been described elsewhere^4^. In this study, we calculated up and down scores based on seven increased genes (*PLBD1, TPST1, MCEMP1, IL1R2, HP, FLT3, S100A12)* and 43 decreased genes (*LCK, CAMK2D, NUP43, SLAMF7, LRRC39, ICOS, CD47, LBH, SH2D1A, CNOT6L, METTL8, ETS1, C2orf27A, P2RY10, TRAT1, BTN3A1, LARP4, TC2N, GPR183, MORC4, STAT4, LPAR6, CPED1, DOCK10, ARHGAP5, HLA-DPA1, BIRC3, GPR174, CD28, UTRN, CD2, HLA-DPB1, ARL4C, BTN3A3, CXCR6, DYNC2LI1, BTN3A2, ITK, SNHG1, CD96, GBP4, S1PR1, NAP1L2, KLF12, IL7R*) of the original 52-gene signature. To study the associations bewteen risk profiles and outcomes, we used one-sided Fisher’s exact test for the discovery cohort, and used chi-squared test for categorical outcomes and two-sample t-test for continuous outcomes (skewed variables were log-transformed) for the validation cohort.

## References

1. Goyal, P., et al. Clinical Characteristics of Covid-19 in New York City. The New England journal of medicine 382, 2372–2374 (2020).

2. Spagnolo, P., et al. Pulmonary fibrosis secondary to COVID-19: a call to arms? Lancet Respir Med 8, 750–752 (2020).

3. Herazo-Maya, J.D., et al. Peripheral blood mononuclear cell gene expression profiles predict poor outcome in idiopathic pulmonary fibrosis. Science translational medicine 5, 205ra136 (2013).

4. Herazo-Maya, J.D., et al. Validation of a 52-gene risk profile for outcome prediction in patients with idiopathic pulmonary fibrosis: an international, multicentre, cohort study. Lancet Respir Med 5, 857–868 (2017).

5. Lee, J.S., et al. Immunophenotyping of COVID-19 and influenza highlights the role of type I interferons in development of severe COVID-19. Sci Immunol 5(2020).

6. Overmyer, K.A., et al. Large-Scale Multi-omic Analysis of COVID-19 Severity. Cell Syst (2020).

7. Wilk, A.J., et al. A single-cell atlas of the peripheral immune response in patients with severe COVID-19. Nature medicine 26, 1070–1076 (2020).

8. Schulte-Schrepping, J., et al. Severe COVID-19 Is Marked by a Dysregulated Myeloid Cell Compartment. Cell 182, 1419–1440 e1423 (2020).

9. Scott, M.K.D., et al. Increased monocyte count as a cellular biomarker for poor outcomes in fibrotic diseases: a retrospective, multicentre cohort study. Lancet Respir Med 7, 497–508 (2019).

10. Chen, Z. & John Wherry, E. T cell responses in patients with COVID-19. Nat Rev Immunol 20, 529–536 (2020).

11. Atyeo, C., et al. Distinct Early Serological Signatures Track with SARS-CoV-2 Survival. Immunity 53, 524–532 e524 (2020).

12. Team, R.C. R: A language and environment for statistical computing. R Foundation for Statistical Computing, Vienna, Austria. (2020).

